# Functional connectivity of fMRI using differential covariance predicts structural connectivity and behavioral reaction times

**DOI:** 10.1101/2021.09.01.458609

**Authors:** Yusi Chen, Qasim Bukhari, Tiger W. Lin, Terrence J. Sejnowski

## Abstract

Recordings from resting state functional magnetic resonance imaging (rs-fMRI) reflect the influence of pathways between brain areas. A wide range of methods have been proposed to measure this functional connectivity (FC), but the lack of “ground truth” has made it difficult to systematically validate them. Most measures of FC produce connectivity estimates that are symmetrical between brain areas. Differential covariance (dCov) is an algorithm for analyzing FC with directed graph edges. Applied to synthetic datasets, dCov-FC was more effective than covariance and partial correlation in reducing false positive connections and more accurately matching the underlying structural connectivity. When we applied dCov-FC to resting state fMRI recordings from the human connectome project (HCP) and anesthetized mice, dCov-FC accurately identified strong cortical connections from diffusion Magnetic Resonance Imaging (dMRI) in individual humans and viral tract tracing in mice. In addition, those HCP subjects whose rs-fMRI were more integrated, as assessed by a graph-theoretic measure, tended to have shorter reaction times in several behavioral tests. Thus, dCov-FC was able to identify anatomically verified connectivity that yielded measures of brain integration causally related to behavior.

## 1 Introduction

Functional connectivity (FC) depends on measures of statistical correlation between brain regions during resting state fMRI recordings (rs-fMRI). The definition of FC has shifted from association to causation, which was previously referred to as effective connectivity [30]. To be consistent with the prevailing definition, we refer to all estimated connectivity patterns, whether causal or not, as FC. Methods for computing FC have been tested on synthetic data, but testing on fMRI recordings is problematic without ground truth. New methods are needed for comparing the performances of these methods using other sources of evidence.

The covariance matrix is the most commonly used method to calculate FC [38]; however, two correlated nodes may not have a direct physical connection because of covariance propagation [43]. Partial covariance reduces indirect correlations but not interactions due to unobserved common inputs. These spurious connections produced by covariance-based methods make it harder to interpret the link between FC and the underlying physiological state. In addition, covariance is a symmetric matrix, which is not consistent with functional dependencies that are not symmetric.

Differential covariance (dCov), which is the focus of this study, reduces false positive connections and recovers the ground truth connectivity in simulation studies [23, 24].^1^ In addition, dCov is, in general, not a symmetric matrix. Unlike other statistical methods, dCov seeks to estimate connections from a dynamical system point of view. It makes use of the derivative signal, in addition to the original signal, to capture the dynamical interactions between neural entities. When recorded neural responses are viewed as a dynamical system governed by a set of ordinary differential equations, the derivative provides information about the moving trend of the state variables. By evaluating the relationship between the derivative signal and the original signal, we can link the current state to future states, and thus extract the directed connectivity pattern. The goal of this paper is to apply dCov to rs-fMRI recordings to evaluate its performance compared with other FC methods in estimating structural connectivity and predicting behavioral data.

Some studies have focused on validating FC on several known projections or subnetworks, such as the default mode network (DMN) [29], but leave open the validity of the larger FC matrix. Since direct activity dependencies between two brain regions depend on fiber projections connecting them, it is reasonable to expect the FC to be closely related to the underlying structural connectivity (SC) [46], with the strength depending on brain dynamics. We used SC obtained through either viral tracing in mice or diffusion Magnetic Resonance Imaging (dMRI) in humans as surrogates for “ground truth”.

The existence of anatomical connections, either direct or indirect, is a necessary condition for the dynamic coupling of neuronal activities between two brain regions. On the other hand, neural activity dependencies are more dynamic and variable than the underlying anatomical linkages. Honey et al. [19] systematically examined the relationship between Pearson correlation based FC and Diffusion Tensor Imaging (DTI) based SC at high spatial resolution. They pointed to the redundancy and unreliability of structurally unconnected functional connections. Since then, major efforts have been made to predict FC from SC [1, 2, 16, 27]. But these approaches typically accounted for less than half of the known structural connections [16], indicating that a complete match from the entire weighted FC matrix to the weighted SC matrix is difficult to achieve owing to limitations of both FC estimation and DTI. To obtain a better match between FC and SC, we chose to focus on examining the subset of significant functional connections. This also had the advantage of increasing the tolerance of FC estimation to various noise sources in fMRI recordings [25].

Another challenge is to gain insights from the estimated FCs into how behavior is generated in brains. Methods from network science, especially those rooted in graph theory, have been useful for analyzing the dynamics of brain communications [3]. Several core concepts describing the segregation and integration of information [40] have provided insights into disease states of the brain [20] and cognitive control mechanisms [26, 35, 7]. This suggests that the topological properties of FCs evaluated by different methods could provide new insights into brain function.

For the above reasons, we systematically benchmarked the link between binarized FC and SC, comparing FC defined by dCov with covariance-based methods and quantifying performance by their correspondence with anatomically strong connections. The FC matrices were also topologically analyzed and compared with behavioral measurements to link them with brain states that give rise to behavior.

## 2 Results

### 2.1 Overview of differential covariance (dCov)

The differential covariance method makes use of the derivative signal. In most neural dynamic models, the derivative of a neuron’s membrane potential is related to its incoming synaptic and intrinsic currents (i.e. sinks), and its own membrane potential is related to its output (i.e., sources) (Fig 1). Differential covariance uncovers the relationship between sinks and sources. In this paper, we calculated three dCov matrices: differential covariance matrix, Δ*c* (Fig S1, Equation 1); partial differential covariance matrix, Δ*p* (Fig S1, Equation 2); and sparse-latent regularized partial differential covariance matrix, Δ*s* (Fig S1, Equation 3).

**Figure 1:**
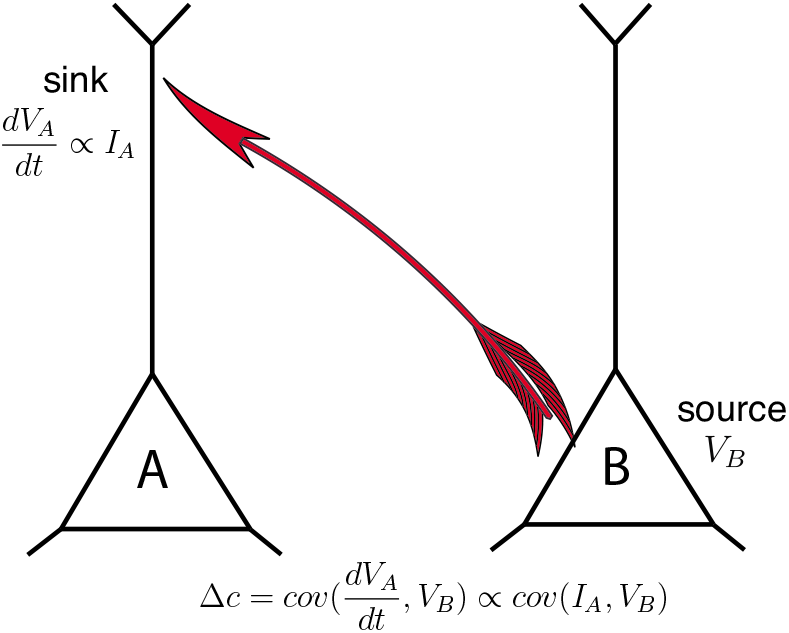
Differential covariance estimates the influence of source activity on changes in the sink activity. The output from the source area is driven by *V*_*B*_ and the input to the sink area is given by 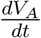, which is related to the input currents, *I*_*A*_. The covariance between these two signals is a measure of the directed flow of information from the sources to the sinks.

First, the (*i, j*) entry of differential covariance Δ*c* is calculated as Equation 1 where *z*_*i*_ is the time trace of node *i* in a network, *dz*_*i*_ is the numerical derivative of *z*_*i*_, the superscript bar denotes the sample mean of a time trace and *cov* denotes the estimation of sample covariance between two time traces.

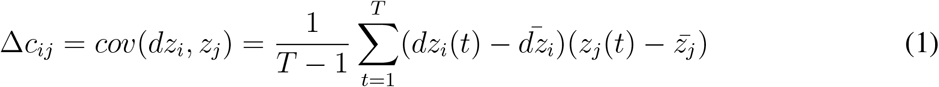

Secondly, the partial differential covariance (Δ*p*) parallels the derivation of the partial covariance matrix [8]. Partial differential covariance controls the effect of confounding variables by calculating the relationship based on residual time traces. In most cases, the residual time traces were obtained by solving a multiple linear regression problem with confounding variables included as independent variables. It follows that the (*i, j*) entry of Δ*p* can be calculated in Equation 2 where *K* is the set of nodes other than *i* and *j*; Cov is the covariance matrix; Cov_*jK*_ ∈ ℝ^1×(*N* −2)^, Cov_*KK*_ ∈ ℝ^(*N* −2)×(*N* −2)^ and Δ*c*_*iK*_ ∈ ℝ^1×(*N* −2)^.

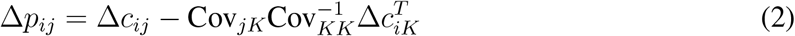

The third matrix Δ*s* was designed to minimize the effects of latent inputs, which are not uncommon in fMRI recordings. Toward this end, we adopted a sparse latent regularization method [48] to separate Δ*p* into an intrinsic connection matrix Δ*s* between observable nodes and a residual matrix *L* representing the influence of latent nodes. In this paper, the intrinsic connection matrix, Δ*s*, was obtained by solving a constrained optimization problem using an augmented Lagrangian multiplier [5]. In Equation 3, Δ*p* is decomposed into a sparse matrix (Δ*s*) and a low-rank matrix (L). The objective function is the weighted sum between L1-norm of the vectorized sparse matrix and the trace of the low-rank matrix. The penalty ratio *α* controls the sparsity of Δ*s*. In this paper, 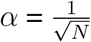.

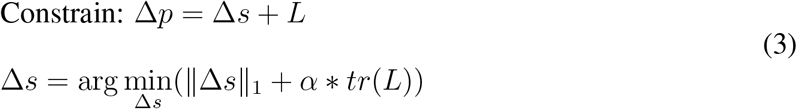

### 2.2 Functional connectivity estimated by differential covariance (dCov-FC)

We followed standard procedures for preprocessing fMRI recordings [36]. The recordings of voxel-wise time traces were decomposed into component-wise time traces and MRI maps of each component through group independent component analysis (ICA) and dual regression (Fig 2A) (Methods). After manually removing artifactual components, the remaining components (30 for mouse and 60 for HCP) were registered with corresponding anatomical locations in a brain atlas (MRI maps are shown in Fig 3 and 4; annotations are shown in Table S2 and Table S3). This pre-processing yielded an *N* ×*T* (number of components × number of time points) matrix representing time-varying haemodynamic signals for various anatomical locations. We then applied a recently developed backward reconstruction method to reconstruct neural signals (*z*) from haemodynamic signals (example time traces shown in Fig S4). Backward reconstruction based on the forward Balloon model [13] has been shown to work well together with dCov applied to synthetic datasets [23].

**Figure 2:**
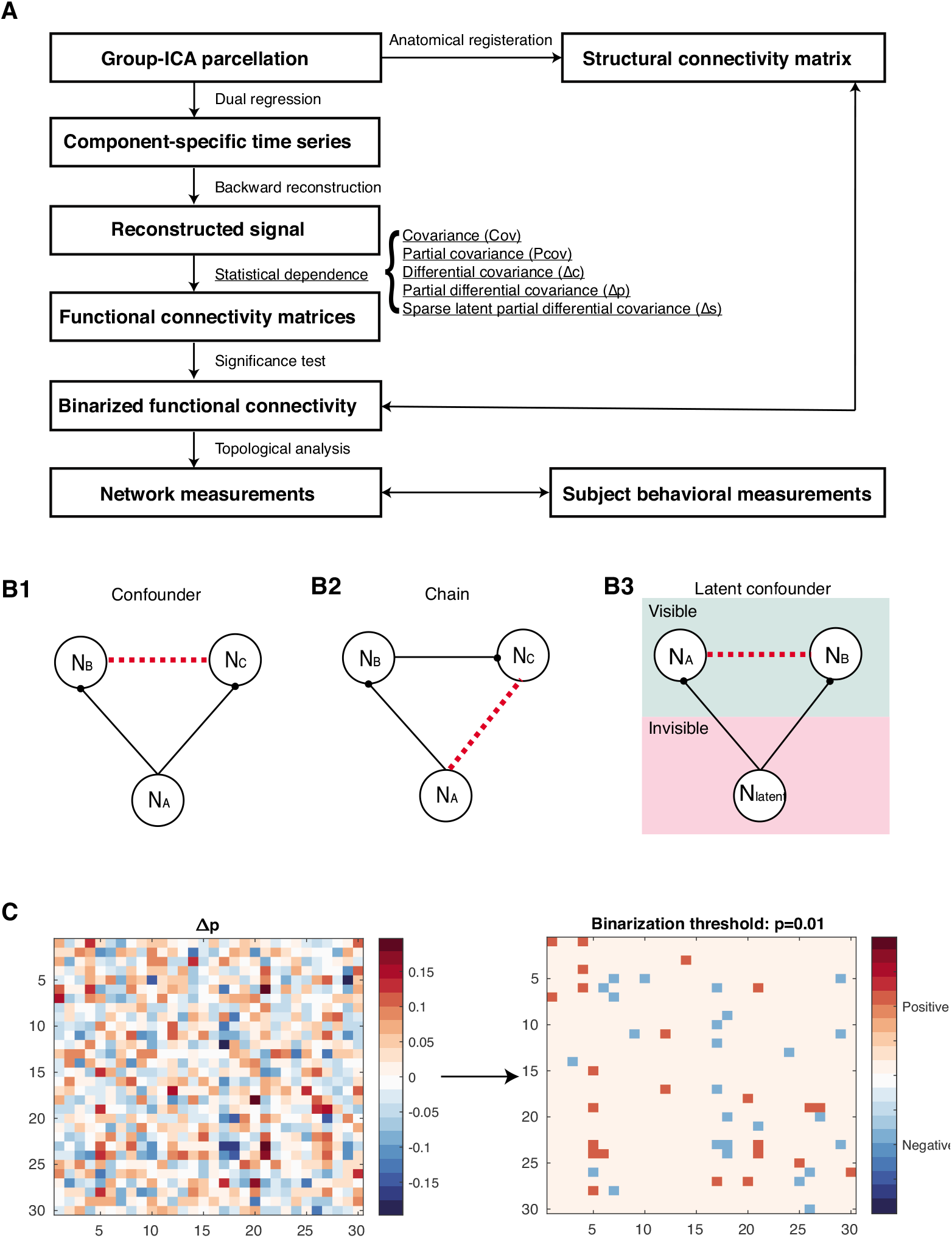
Workflow and features of differential covariance based methods. A) Workflow. B) Differential covariance based methods have been shown to effectively reduce the three types of errors in network modeling shown in B1, B2 and B3 [24]. Black solid lines denote actual physical connections and red dashed lines denote false positive connections that are absent. B1: confounder effects due to common input; B2: chain effects due to propagation; B3: latent confounders due to unobserved common inputs; C) The partial differential covariance matrix, Δ*p*, on the left and its binarized matrix on the right (binarization threshold = 0.01) of one mouse subject. Partial differential covariance matrices are directed, nonsymmetric and sparser than the corresponding covariance matrices (Fig S1).

**Figure 3:**
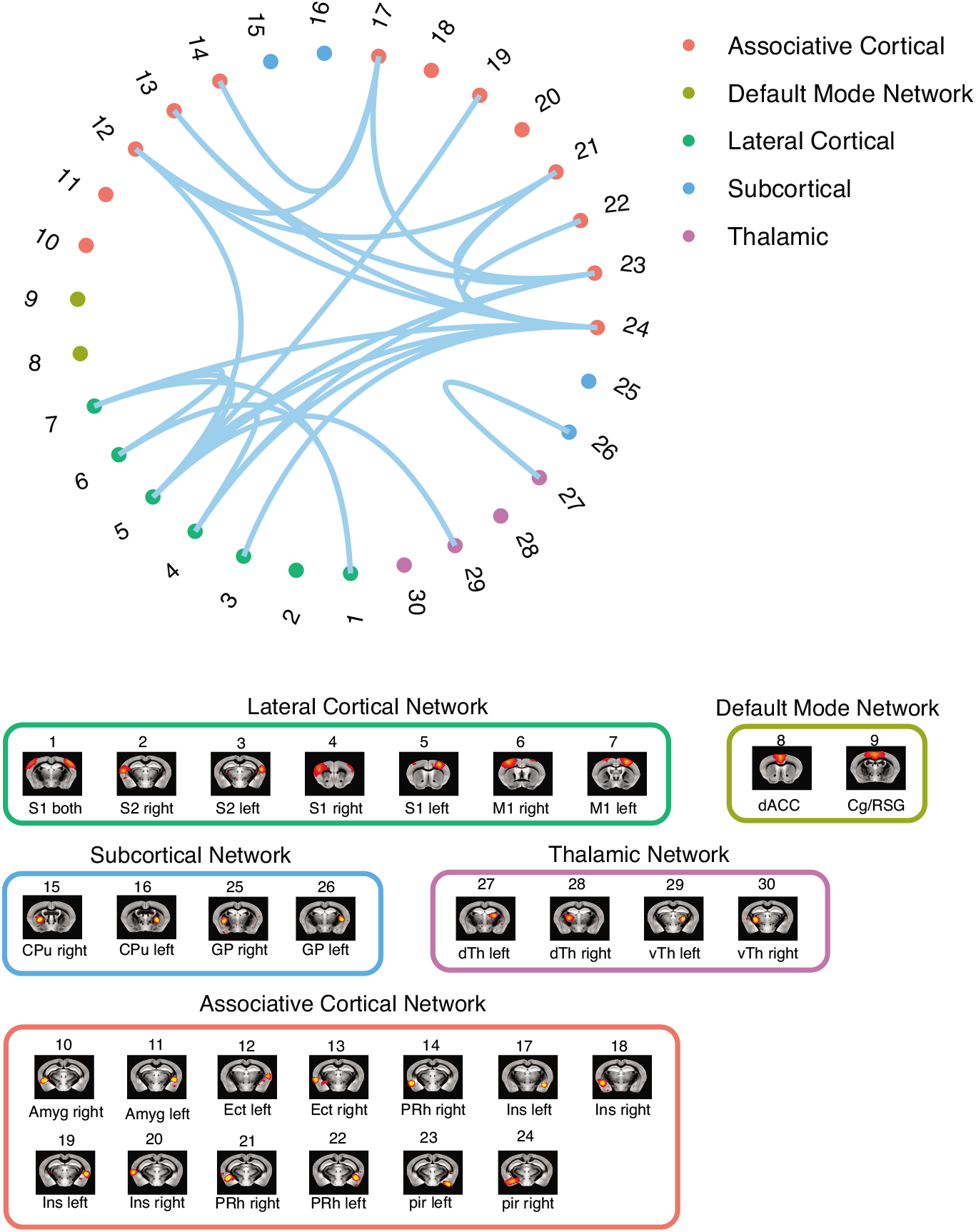
Significant Δ*p*-FC connections shared across subjects and associated MRI spatial maps from the mouse rs-fMRI dataset. (A) For Δ*p*-FC estimated from the mouse dataset, connections which are significant (binarization threshold = 0.01) in more than 2 subjects (out of 12) were plot in the circular plot; (B) MRI spatial maps of the independent components were grouped into sub-networks. The registered anatomical locations were shown underneath. Note that there could be more than one component (for example, component 17 and 19) corresponding to the same anatomical region due to the unsupervised nature of group ICA. S1: primary somatosensory cortex, S2: secondary somatosensory cortex, M1: primary motor cortex, dACC: dorsal anterior cingulate cortex, Cg: cingulate cortex, RSG: retrosplenial cortex, Amy: amygdalar, Ect: entorhinal cortex, PRh: perirhinal, CPu: caudoputamen, Ins: insular, pir: piriform, GP: Globus Pallidus, dTh: dorsal thalamus; vTh: ventral thalamus. Refer to Supplementary Table S3 for detailed annotation.

**Figure 4:**
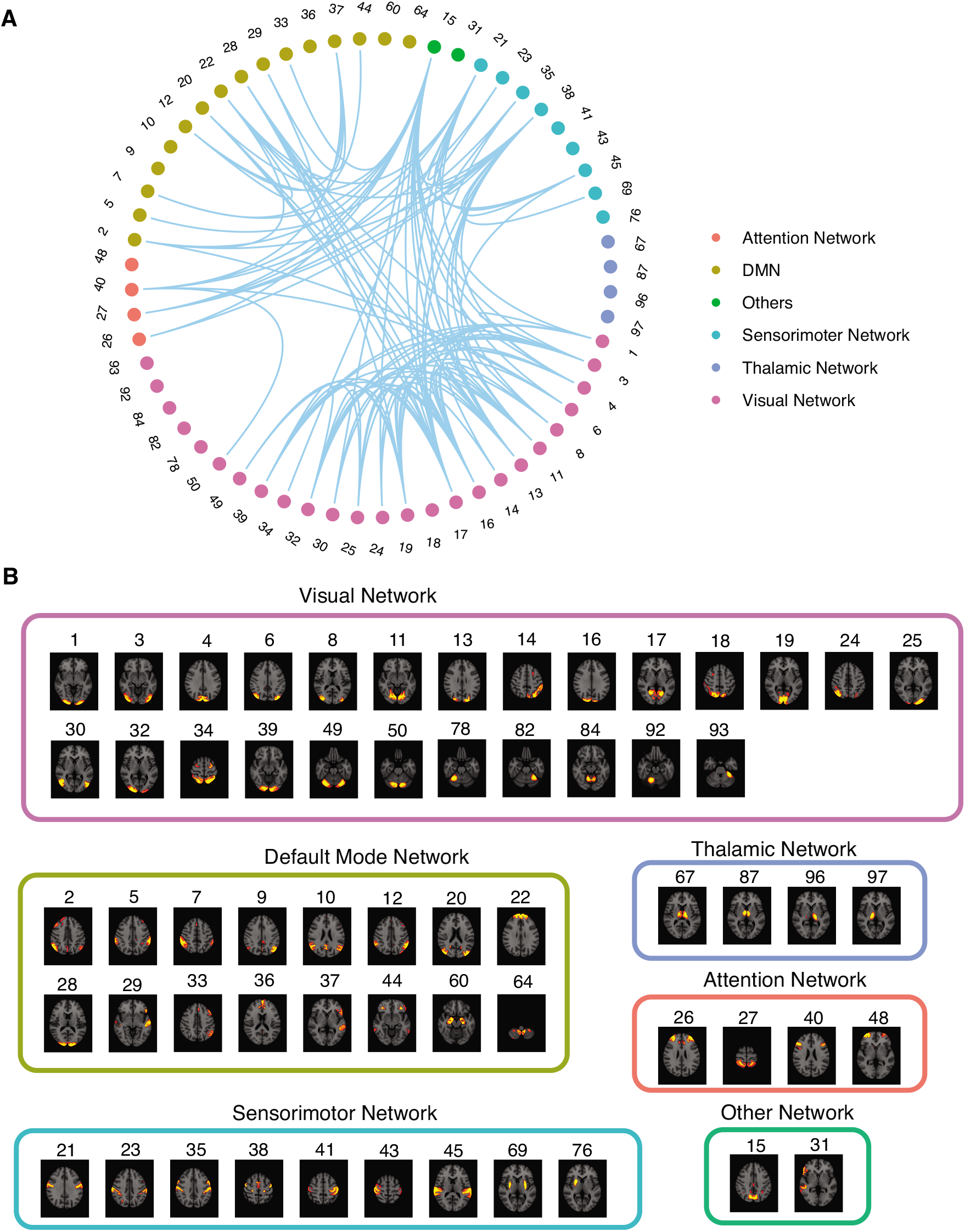
Significant Δ*p*-FC connections shared across subjects and associated MRI spatial maps from the HCP rs-fMRI dataset. (A) For Δ*p*-FC estimated from the HCP dataset, connections which are significant (binarization threshold = 0.01) in more than 30 subjects (out of 1003) were plot in the circular plot; (B) MRI spatial maps of the independent components were grouped into sub-networks. The number marks the index of the independent components. The indices are from 1 to 100 but only 60 of them were shown here because the remaining components were treated as imaging artifacts. Refer to Supplementary Table S2 for more detailed annotation.

Functional connectivity (FC) matrices (*N* × *N*) were estimated from the reconstructed neural signals, *z*, using either covariance-based methods, including the covariance matrix and the partial correlation matrix, or the three dCov matrices mentioned above. Differential covariance has been shown to specifically reduce three types of common false positive connections (Fig 2B). To be more specific, Δ*c* dealt with errors due to confounder effects (Fig 2B1). In the calculation of Δ*p*, both confounder and chain effects (Fig 2B2) were taken into account. Finally, a sparse latent regularization was used in the calculation of Δ*s* (Equation 3) to reduce the effect of latent confounders assuming this latent effect is low-rank (Fig 2B3). Note that in general, dCov matrices give rise to FC matrices that are not symmetric (antisymmetric for Δ*c*), which could, in practice, potentially point to the direction of information (Fig S1).

To reduce the susceptibility of FC matrices to noise, we determined the significance level of connections using a bootstrap procedure and the null hypothesis is that activities of every node in the network are generated independently. An autoregressive (AR) bootstrap procedure [9, 28] was used to generate the null time traces to preserve the power spectral density of haemodynamic signals, and then null connection matrices were estimated from the null time traces. For each empirical connection, the probability that its value belonged to the null distribution, its *p* value, was calculated assuming the distribution was Gaussian. Thresholds of significance were applied to binarize the FC matrix so that we could focus on the set of significant connections. Because dCov-based methods intrinsically reduce false positive connections, they yielded fewer significant connections compared to covariance-based methods (Fig S1)

We applied the above workflow to extract covariance-based and dCov FC matrices from fMRI recordings in anesthetized mice and resting-state recordings from the HCP dataset. The mouse dataset included 12 subjects, and the HCP dataset included 1,003 young adult subjects. Fig 3B and 4B show the MRI spatial maps of each component. Due to the individual variability of connectivity patterns, the circular plots in Fig 3A and 4A only showed the subset of significant connections shared across subjects. Components were allocated into networks according to their corresponding anatomical locations. The first inspection of shared connections showed promising correspondence between Δ*p*-FC and the anatomical wiring of different brain regions. For example, in Fig 3, Δ*p*-FC picked up important connections within the mouse lateral cortical network, which is located in the densely interconnected M2 module reported by Swanson et al. [44]. In Fig 4, the visual network is densely interconnected across multiple subjects revealed by Δ*p*-FC. Therefore, we next quantified the similarity between dCov-FC and SC at individual level and population level in both human and mouse subjects.

### 2.3 Differential covariance (dCov) FC more closely matches the underlying structural connections

We developed a pipeline to systematically quantify the similarity between FC and SC. We first constructed an *N* × *N* structural connectivity, “ground truth” matrix, with nodes corresponding to the anatomical regions that matched the independent components obtained from resting-state recordings (Fig S2 and S3, Methods). We used the existing SC database for the connectivity profiles between different anatomical regions. Specifically, we used population level viral tracing data [44] for mouse SC and individual level dMRI data [32] for human SC (Methods). The mouse “ground truth” matrix (Fig S2) is not symmetric while the human “ground truth” matrices (Fig S3) are symmetric and sparse with substantial amount of individual variability. In both cases, higher values indicate higher structural connectivity strength. We then calculated the average structural connectivity strength (ASCS) of the set of significant connections picked through AR bootstrapping under multiple binarization thresholds. The decreasing binarization threshold provides a stricter criterion for connection selection and pictures an asymptotic description of the evaluated quantity. A higher ASCS value implies that a specific FC more closely matched the underlying SC.

We calculated ASCS values for FC defined by the covariance matrix (cov), partial correlation matrix (Pcov), Δ*c*, Δ*p* and Δ*s*. Fig 5B and Fig 5D show ASCS values of significant connections, using multiple binarization thresholds, pooled from all subjects in mouse and HCP dataset, respectively. In both mouse (Fig 5B) and human datasets (Fig 5D), Δ*p* and Δ*s* had significantly (p<0.05, rank-sum test) higher ASCS values than those based on the covariance matrix. Fig 5A and Fig 5C showing structural connectivity strength (SCS) distributions from one individual subject revealed a reduced number of low-SCS connections in both Δ*p* and Δ*s*. This is a consequence of the method’s ability to reduce false positive connections, as previously shown using synthetic data in simulation studies [23].

**Figure 5:**
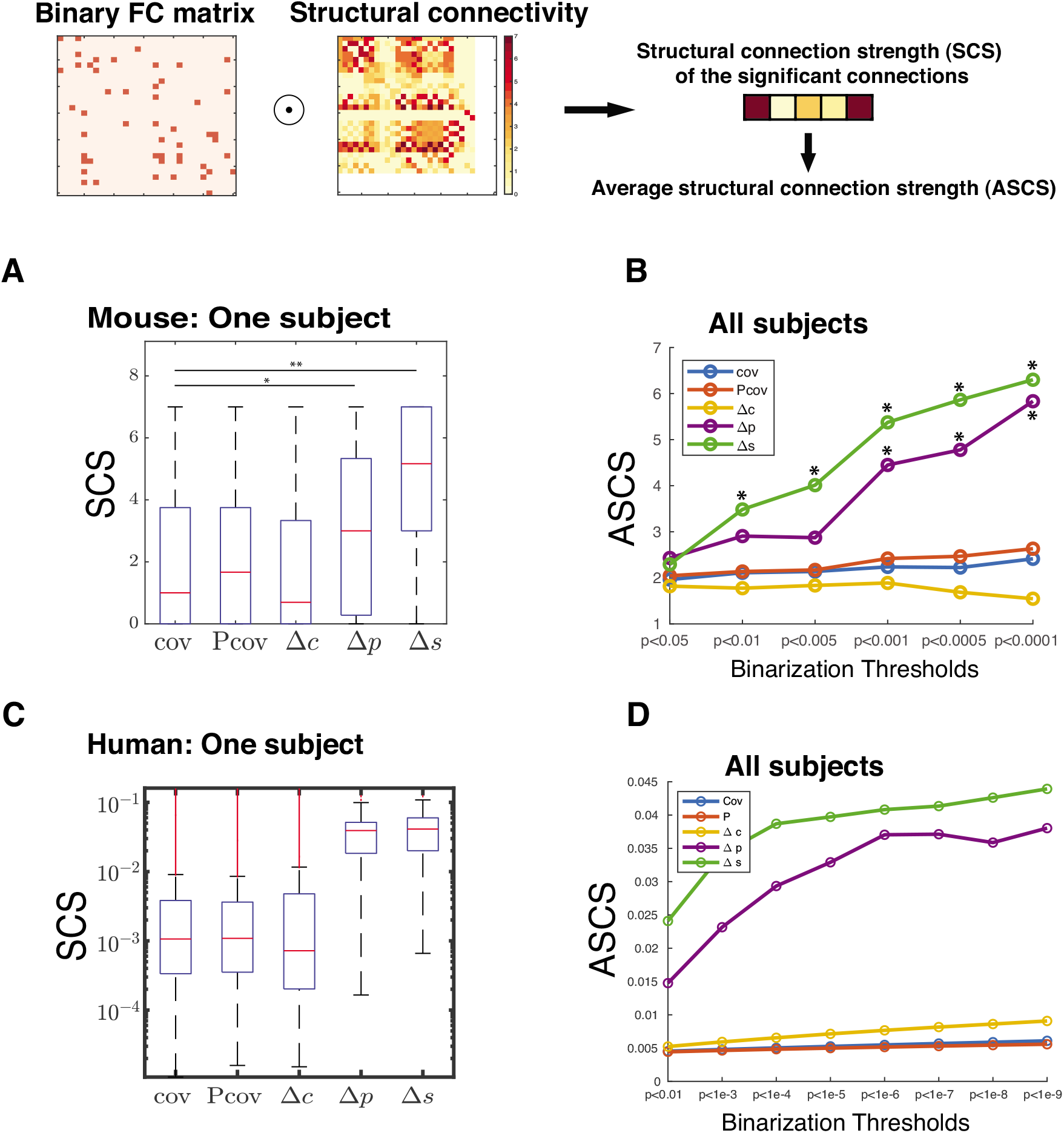
Differential covariance (dCov) based FC preferentially identified connections with high SCS in both mouse and HCP dataset. A) C) The SCS distribution of the significant (p<0.01, rank-sum) connections identified by different methods from recordings of one mouse subject (A) or one human subject in the HCP dataset (C). Δ*p* and Δ*s* identified connections with significantly higher SCS. B) D) Under different binarization thresholds, ASCS of the significant connections pooled from all subjects in the mouse dataset (B) and the HCP dataset (D). Asterisk mark implies a significant difference (p<0.01, rank-sum) compared to the quantity identified by the covariance method. Under most of the binarization thresholds, Δ*p* and Δ*s* have better performance than the baseline covariance method in picking up structurally connected regions. Abbreviations: FC: functional connectivity; cov: covariance; Pcov: partial covariance; Δ*c*: differential covariance; Δ*p*: partial differential covariance; Δ*s*: partial differential covariance with sparse latent regularization.

### 2.4 Differential covariance (dCov) FC measures the integration of brain activity

Next, we wanted to know whether dCov based FC matrices, which are aligned with the underlying SC, could also provide insights into the interpretation of behavioral measurements. The HCP dataset (Methods) consists of not only high-quality macroscopic-level imaging data, but also hundreds of behavioral measurements, including detailed psychological test results from 1003 young adult subjects [45]. The large sample size and the full assessment of subjects’ psychological states render the HCP dataset an invaluable source for investigating the organizing principles linking human brain networks and behavioral readouts.

We calculated several topological properties of the FC matrices [33]. Because of the inherent interdependencies and transitivity of bivariate methods, the covariance matrix is not suitable for calculating network properties [3]. So FC derived from the partial correlation matrix (Pcov-FC) was used for control. Pcov-FC and Δ*s*-FC exhibited similar network properties such as network degree distribution (Fig S5A), clustering behavior, transitivity and modularity properties (Fig S5C) across subjects. In addition, both Pcov-FC and Δ*s*-FC showed similar network degree distribution to that of population level diffusion tensor imaging network reported in Hagmann et al [18] (Fig S5B). Surprisingly, the network efficiency of Δ*s*-FC is strongly anti-correlated (r=0.71, p=4.8e-152) with that of Pcov-FC (Fig 6A). Network efficiency, calculated as the average of the inverse path length of shortest signaling routes between two nodes (Methods), measures the network’s integration level. Since Δ*s*-FC reduces false positive connections/paths in the network, it selects for the important and reliable routes and decreases redundant routes. As a consequence, network efficiency of Δ*s*-FC could potentially be a better description of the integration level.

**Figure 6:**
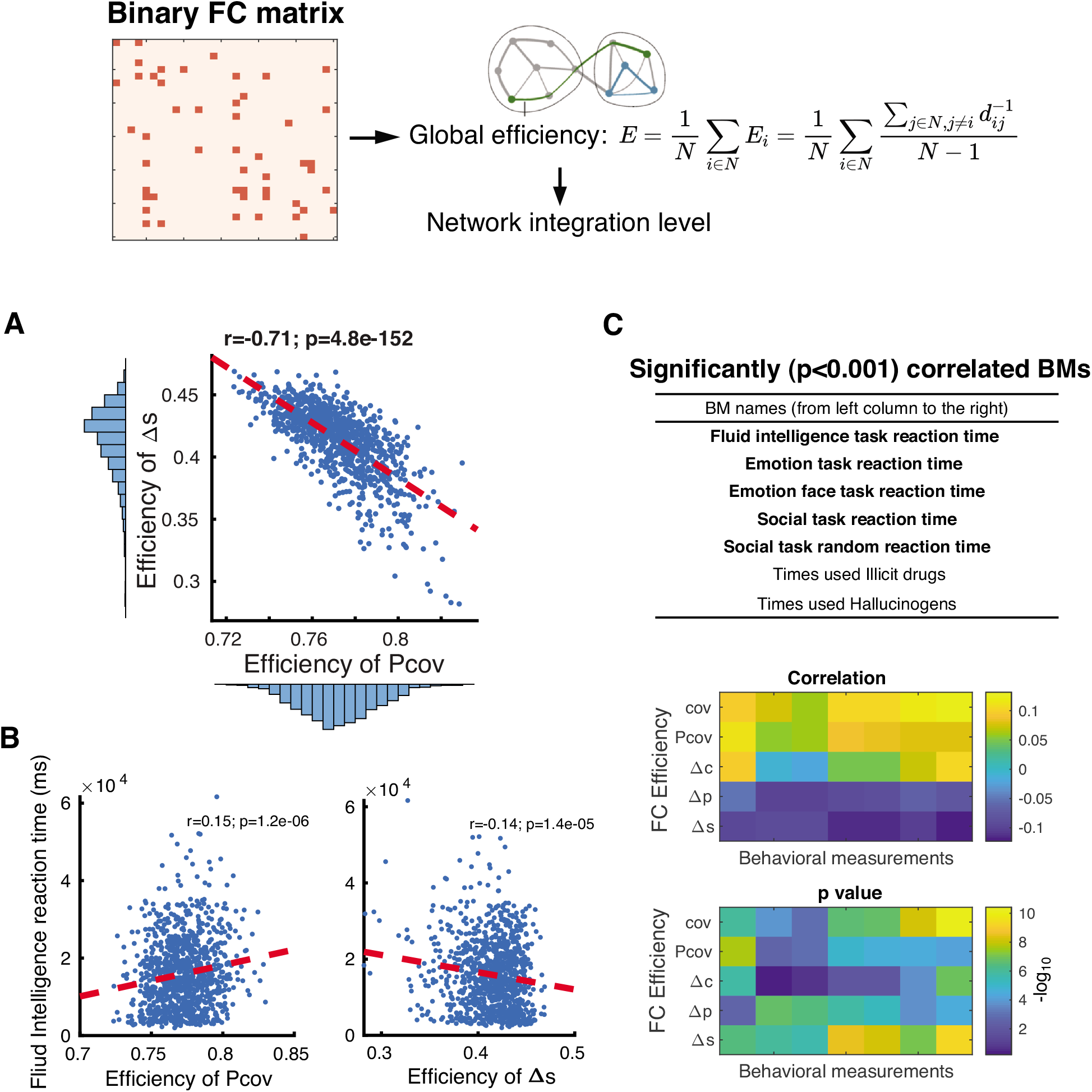
Differential covariance based FC matrices could be a more reasonable measurement of the integration level of the brain. A) In the HCP dataset, global efficiency of Δ*s*-FC and Pcov-FC are strongly anti-correlated (r=0.71, p=4.8e-152). B) Global efficiency of Pcov-FC is positively correlated (left, r=0.15, p=1.2e-6) to the reaction time of fluid intelligence task while the global efficiency of Δ*s*-FC is negatively correlated (right, r=−0.14, p=1.4e-5) to it. In most cases, we would expect a negative correlation because the more highly integrated the network, the shorter the response time. C) The table lists the behavioral measurements that are most significantly correlated (p<0.001) to the global efficiency of Δ*s*-FC, after controlling for confounding factors. Most of the significant behavioral measurements are related to reaction time in different tasks (shown in bold). Heat maps below show their correlation values and p values.

To investigate this hypothesis, we compared this measure of network integration with each subject’s fluid intelligence quantified by their reaction time to the test. This behavioral measurement has been previously shown to be most related to FC [39]. One expects that a higher level of network integration would lead to shorter reaction times, and indeed Δ*s*-FC based global efficiency is inversely correlated (r=−0.14, p=1.4e-5) to reaction time (Fig 6B). As a control, we sorted the correlations between Δ*s*-FC based global efficiency and all behavioral measurements after controlling common influencing factors like age and brain size. Five out of seven significant (p<0.005) behavioral measurements are reaction time in various psychological tests (Fig 6C), which further confirmed the significance of Δ*s*-FC as a measure of the integration of brain activity.

### 3 Discussion

In this paper, we applied three variants of dCov to estimate FC from resting-state fMRI recordings, Δ*c*, Δ*p* and Δ*s*. Compared to the covariance matrix, differential covariance identified a subset of connections with high structural connectivity strength, largely owing to dCov’s ability to reduce three types of false positive connections. These results were confirmed in both human and mouse rs-fMRI recordings. In addition, by ruling out redundant signaling routes, the sparse-latent regularized partial differential covariance matrix (Δ*s*-FC) provided the closest link to behavioral measurements and functional network integration based on the inverse of shortest path length, a measure of global efficiency.

The estimates of directed connectivity afforded by dCov-FC are formally related to the estimates of effective connectivity from dynamic causal modeling, under linear assumptions, because dCov-FC rests upon the same generative model used in linearized dynamic causal models [11]. However, interactions are estimated using variational Bayesian inference in dynamic causal modeling [12], whereas dCov-FC expresses the interaction pattern in terms of the covariance between the derivative signal and the original signal. Without the need to go through the model fitting process, dCov is a much faster and more straightforward algorithm.

The current versions of dCov assume that the underlying system is linear and that the connectivity pattern is closely related to the transition matrix. Although the computational processes in brains are by no means linear, the averaging of activity in large neural populations, as in the case of fMRI recordings, could be quite linear [41]. The current linear dCov estimator could be extended to nonlinear estimators, taking into account the threshold firing properties of individual neurons, to better extract connectivity at the single neuron level.

The calculation of sparse-latent regularized partial differential covariance matrix (Δ*s*) used a method for sparse-latent regularization that was initially proposed by Yatsenko et al. [48] and Candes et al. [5]. This sparse-latent regularization assumes a sparse structure of the intrinsic connections and a low-rank component representing common fluctuations and common latent inputs. This assumption may or may not be fulfilled in fMRI recordings, and there is no straightforward way to test this assumption. Nonetheless, the addition of sparse latent regularization did not influence the qualitative conclusions (compare Δ*p* and Δ*s* in Fig 5 and Fig 6C). In addition, tractography methods are known to underestimate long-range connections, and thus dMRI-based SC may bias towards a local and sparse connectivity pattern [21].

Another concern is that the correspondence between dMRI based SC and dCov-FC could be due to the fact that both methods favor sparse configurations. However, the sparse structure doesn’t necessarily guarantee a higher chance of sharing nonzero entries. In addition, the mouse SC matrix (Fig S2A) constructed by viral tracing is not sparse, but dCov-FC was nonetheless better able to identify the structurally strong connections.

In conclusion, in addition to validating dCov’s performance on fMRI recordings, we have also extended previous studies linking FC to connectivity and behavior. The analytic techniques for studying links to anatomy and behavior introduced here can be applied to other methods for determining FC and other datasets, which could provide stronger confidence in their interpretation.

## 4 Methods

### 4.1 Data acquisition and pre-processing

We used two fMRI datasets, including an anesthetized mouse dataset first reported in Bukhari et al [4] and the extensively processed young adult resting state fRMI (rs-fMRI) S1200 dataset from Human Connectome Project (HCP, https://www.humanconnectome.org).

For the mouse dataset, experiment and imaging protocols were defined in the original publication. For all 12 animals, each subject received approximately 1.2% Isoflurane (Abbott, Cham, Switzerland) with a tolerance of 0.1% in a 20% oxygen/80% air mixture. The data was collected in two sessions, where the animals remained inside the scanner during the whole time. The total time series acquired during the two sessions was of 720s (680s) length. Subjects were scanned using a Bruker Biospec 94/30 small animal MR system (Bruker BioSpin MRI, Ettlingen, Germany) operating at 400 MHz (9.4 T). A gradient-echo echo-planar imaging (GE-EPI) sequence has been used for rs-fMRI data acquisition with field of view = 16 × 7 mm^2^, matrix dimensions = 80 × 35, TR = 1 second, TE = 12 ms, flip angle = 60 degree. Each subject had 680 imaging frames collected.

All the pre-processing was performed using tools from FMRIB’s Software Library (FSL version 5). FSL’s recommended pre-processing pipeline was used. Motion correction, removal of non-brain structures, high pass temporal filtering with sigma = 75.0 s; pre-whitening and global spatial smoothing using a 0.2 mm Gaussian kernel was applied to increase signal to noise ratio as part of the pre-processing. After the pre-processing, the functional scans were aligned to the high-resolution anatomical AMBMC template (http://www.imaging.org.au/AMBMC/AMBMC) using linear affine and nonlinear diffeomorphic transformation registration as implemented in ANTs (ANTs. v 1.9; http://picsl.upenn.edu/ANTS/). We used FSL’s MELODIC for probabilistic independent component analysis. The multi-session temporal ICA concatenated approach, as recommended for rs-fMRI data analysis, allowed to input all subjects from all the groups in a temporally concatenated fashion for the ICA analysis. ConcatICA yielded different activations and artifact components without the need of specifying any explicit time series model. A total of 100 independent components (IC maps) were extracted and the mixture model approach was applied on these estimated maps to perform inference analysis. An alternative hypothesis test based on fitting a Gaussian/gamma mixture model to the distribution of voxel intensities within spatial maps 53,54 was used to threshold the IC maps. A threshold of 0.5 (p < 0.5) was selected for the alternative hypothesis in order to assign equal ‘cost’ to false-positives and false-negatives. Out of the 100 independent components (IC maps), only 30 numbers of components were selected, while the components that overlapped with vascular structures and ventricles were excluded from further analysis. Similarly, regions at the brain surface, which are prone to be affected by the motion artifacts due to e.g. breathing, were excluded.

For the HCP dataset, we used the extensively processed “HCP1200 Parcellation + Timeseries + Netmats (1003 Subjects)” dataset available through the website. Detailed pre-processing and study design could be easily accessed through the website. In this release, 1003 healthy adult human subjects (ages 22-37 years, 534 females) were scanned on a 3-T Siemens connectome-Skyra scanner (customized to achieve 100 mT m^−1^ gradient strength). Each subject underwent 4 × 15 minutes recording sessions with temporal resolution of 0.73 second and spatial resolution of 2 mm isotropic. Subject-specific measures including behavioral, demographic, psychometric measures are available from the HCP data dictionary.

For imaging data processing, each 15-minute run of each subject’s rs-fMRI data was pre-processed according to Smith et al [36]; it was minimally-preprocessed [15], and had artefacts removed using ICA+FIX [34] [17]. Inter-subject registration of cerebral cortex was carried out using areal-feature-based alignment and the Multimodal Surface Matching algorithm (‘MSMAll’) [31] [14]. Each dataset was temporally demeaned and had variance normalization and then fed into the MIGP algorithm, whose output is the top 4500 weighted spatial eigenvectors from a group-averaged PCA (a very close approximation to concatenating all subjects’ timeseries and then applying PCA) [37]. The MIGP output was fed into group-ICA using FSL’s MELODIC tool, applying at several different dimensionalities (D = 25, 50, 100, 200, 300). In our analysis, we used the 100-dimension decomposition.

### 4.2 Component specific time series

For a given “parcellation” (group-ICA map), the set of ICA spatial maps was mapped onto each subject’s rs-fMRI time series data to derive one representative time series per ICA component per subject. This process was fulfilled by the standard “dual-regression stage-1” approach, in which the full set of ICA maps was used as spatial regressors against the full data [10]. This results in an *N* × *T* (number of components × number of time points) matrix for each subject. Thus, we consider each ICA component as a network “node”. These time series can then be used to reveal statistical relationship in a network, as described in Section 4.4.

### 4.3 Reconstruction of neural activity

Before estimating the functional connectivity, we adopted two methods to reconstruct neural signals *z* from haemodynamic time series *y*, which was obtained from the dual regression process described above.

#### 4.3.1 Backward reconstruction

Backward reconstruction was developed based on a biophysical model which transforms neural activities to BOLD signals via a set of differential equations[42]. One important step in this model links blood flow to BOLD signal through a Balloon model assumption. Therefore, in this paper, we will refer to this entire biophysical process as the forward Balloon model. The backward reconstruction process linearizes the equations and derives a surrogate neural signal *z* as a linear combination of haemodynamic signals *y* and its *i*^*th*^ order derivatives *y*^(*i*)^ (Eq. 4) [22]. Note that according to the linearized Balloon model, the reconstructed surrogate signal *z* is a linear combination of real neural signal *V* and its first order derivative 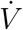,

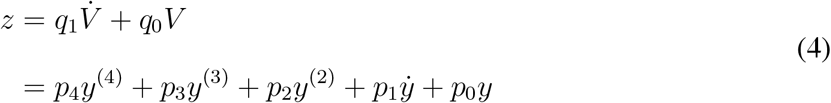

The coefficients *p*_*i*_ and *q*_*i*_ are functions of biophysical parameters: *τ*, *ϵ*, *α*, *κ*, *γ*, *ϑ*_0_, *ρ*, *η* and *V*_0_ (Eq. 5). Some parameter values vary with respect to magnetic field strength (B). Please refer to Table S1 for annotations of these biophysical parameters and the specific values used in this paper. Most values were taken from Stephan et al [42] unless cited. We also demonstrated that the backward reconstruction process is robust to a wide range of parameter values in simulation tests (data not shown).

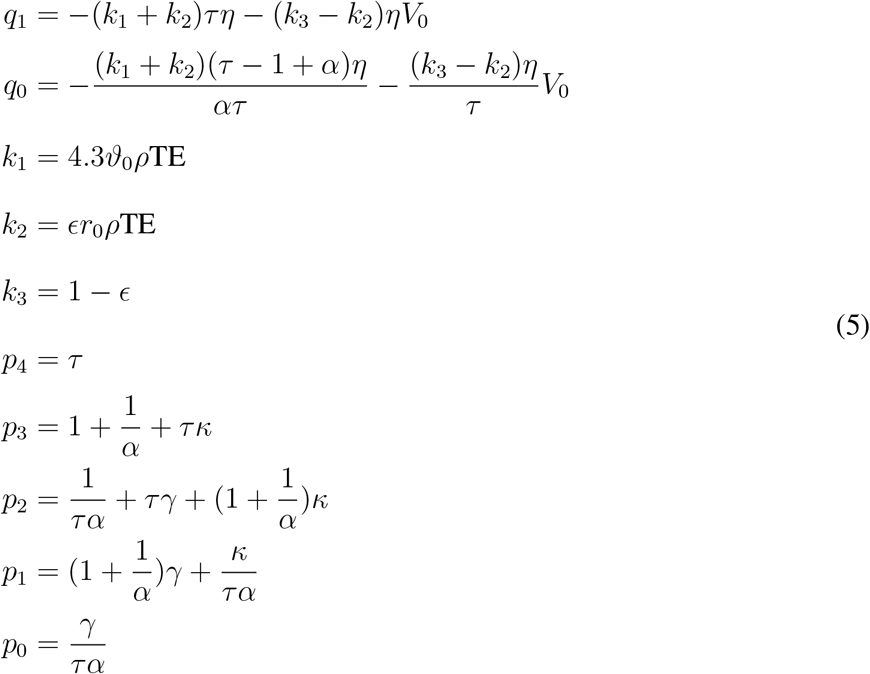

#### 4.3.2 Blind deconvolution

Blind deconvolution is an alternative method to remove effects of hemodynamic response from rs-fMRI recordings [47]. The basic idea is to consider the fluctuations of rs-fMRI as ‘spontaneous event-related’, individuate point processes corresponding to signal fluctuations with a given signature, extract a region-specific Hemodynamic Response Function (HRF) and use it for deconvolution. Details and implementation could be found in the original paper [47].

### 4.4 Estimation of Functional Connectivity

Functional connectivity is represented by a *N* × *N* matrix which describes the statistical relationship of *N* node-time series. To quantify this relationship, we adopted two commonly used covariance based algorithms and three novel differential covariance based algorithms.

#### 4.4.1 Covariance based Methods

Denote the entire *N* node-time series as a *N* × *T* matrix with each row vector as *z*_*i*_, then entry (*i, j*) of the covariance matrix Cov ∈ ℝ^*N* ×*N*^ could be calculated as follows:

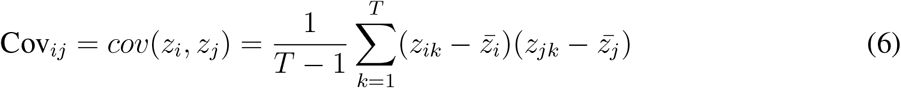

where the superscript bar denotes the sample mean of a time trace and *cov* denotes the estimation of sample covariance between two time traces.

The partial covariance matrix Pcov ∈ ℝ^*N* ×*N*^, also called the precision matrix, aims to estimate direct connections more accurately than the full covariance matrix. Pcov is the inverse of the covariance matrix:

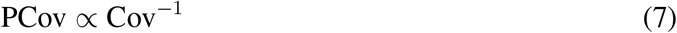

#### 4.4.2 Differential Covariance Based Methods

Differential covariance based methods are a set of algorithms developed to extract direct connections in a network, especially under the influence of common inputs and latent inputs. The calculation of dCov based FC was detailed in Section 2.1 of main text. In our previous work [24][**?**], the advantages of differential covariance based methods over covariance based methods have been fully demonstrated both analytically and numerically. Readers can refer to our previous work for more details.

### 4.5 Significance Testing of FC

To assess the statistical significance of the relationship between hemodynamic signals, we used an autoregressive (AR) bootstrap procedure [9][28] to preserve the power spectrum density (PSD) of hemodynamic signals. For a specific connection, denoted as element (*i, j*) of the above FC matrices, our null hypothesis was that BOLD signal *z*_*i*_ and *z*_*j*_ were not statistically related regardless of other nodes’ time traces. To generate null time series, we fit separate AR processes of model order *q* to node-specific time traces. The model order *q* was determined according to the Bayesian information criterion (BIC). A higher order model was rejected if it could not decrease BIC by more than 2. Using the estimated AR coefficients of empirical time series, we generated 1000 surrogate null time series and then computed the associated functional connectivity corresponding to the null hypothesis. For each connection, we assumed a Gaussian distribution of the null connectivity values generated from null time traces. P value was calculated as the probability of the empirical FC value appeared under the null Gaussian distribution. In this paper, we adopted different significance levels as binarization thresholds.

### 4.6 Structural Connectivity (SC) Matrix

Structural connectivity matrix was constructed as an *N* × *N* matrix with each node corresponding to the anatomical regions represented by one IC map.

To construct the structural connectivity matrix for mouse subjects, we first manually identified the anatomical regions shown on each IC map and then pulled out their connectivity profiles from existing SC database. If one IC map corresponds to multiple anatomical locations in the database, following the method in Hagmann et al [18], we calculated the average connectivity strength of these locations as the connectivity strength of this IC map. The adopted SC database quantified axonal connections within rat cerebral cortex and nuclei [44]. Connection strength was quantified from 1=lowest to 7=highest. The database was built upon a systematic review of neuro-anatomical literature reporting tracing results. The diagonal of mouse SC matrix was set to 7 because intra-regional connections are intensive and there is limited literature reporting intra-regional tracing.

For the structural connectivity of human subjects, we used the individual level SC matrices constructed in our previous work [6] (Refer to method 4.3 in the reference). The mapping between ICs and dMRI measurements were performed at voxel level. Only 46 HCP components were used in the FC-SC analysis since dMRI measurements were restricted to cortical voxels. Readers could refer to Table S3 (mouse) and Table S2 (HCP) for the identified anatomical regions of each IC map.

### 4.7 Network Topological Analysis

For the HCP dataset, the estimated FC matrices were binarized (threshold=0.05) and then assessed using topological network measurements so that we could characterize the FC network using several neuro-biologically meaningful and easily computable measurements [33]. These measurements include node degree distribution (*k*_*i*_), global efficiency (*E*), clustering coefficient (*C*), transitivity (*T*), local efficiency (*E*_*loc*_) and modularity (*Q*). All the measurements were computed using brain connectome toolbox (https://sites.google.com/site/bctnet/). For a binarized network with *N* nodes, these quantities were calculated as follows:

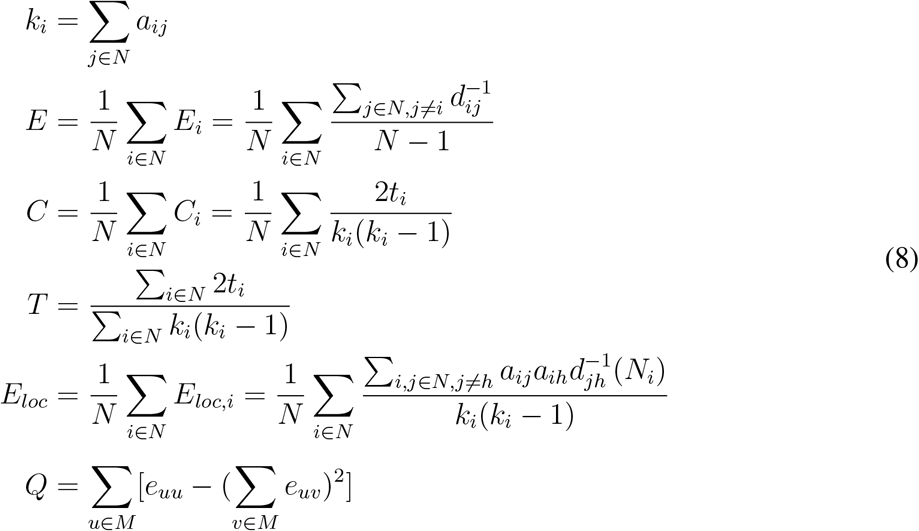

where *a*_*ij*_ ∈ {0, 1} refers to the existence of network edges, *d_ij_* refers to shortest path length between two nodes (*d*_*ij*_ = ∞ if not connected), *t_i_* is the number of local triangles around node *i*. In the calculation of modularity (*Q*), the network was fully subdivided into a set of nonoverlapping modules *M* and *e_uv_* is the proportion of all links that connect nodes in module *u* with nodes in module *v*.

To compare the topological properties of different FC matrices, we calculated the correlation *E*, *C*, *T*, *E_loc_* and *Q* between Pcov-FC and Δ*s*-FC across all subjects (Figure S5C). The node distribution of Pcov-FC, Δ*s*-FC from an example subject and Diffusion Tensor Imaging (DTI) [18] network was fitted by Gaussian distribution, power law distribution (small value truncated) and exponential distribution using maximum likelihood estimation (Figure S5A).

To evaluate the relationship between global network efficiency (*E*) and behavioral measurements (BM), we first combine all BMs documented in public dataset and restricted dataset (all available to download from the HCP website), resulting a matrix of 1003 × 408 (number of subjects × number of SMs). Then in reference to Smith et al [39], we identified five confounding BMs, including age, weight, height, blood pressure-systolic, blood pressure-diastolic and total brain volume. All these confounding BMs were regressed out of the network efficiency measurements *E* so that we only focused on the residual efficiency values. We further excluded BMs with more than 200 missing data points. The Pearson correlation between residual efficiency and BMs are calculated and BMs are sorted according to the significance of their correlation between Δ*s*-FC (Figure 6C).

### 4.8 Data and code availability

The MATLAB code used to run the analysis is available at https://github.com/yschen13/DiffCov_fMRI. The HCP dataset is publicly available at https://www.humanconnectome.org.

## Acknowledgement

We thank Dr. Thomas Liu for helpful discussions and for suggesting to look for behavioral correlates of dCov. The authors would like to thank Burke Rosen, Javier How, Robert Kim, Ben Tsuda and other CNL members for helpful discussion and feedback. The authors also thank Jorge Aldana for assistance with computing resources. This research was supported by the Office of Naval Research (N00014-16-1-2829) and NIH/NIBIB (R01EB026899). Human rs-fMRI dataset was provided by the Human Connectome Project, WU-Minn Consortium (Principal Investigators: David Van Essen and Kamil Ugurbil; 1U54MH091657) funded by the 16 NIH Institutes and Centers that support the NIH Blueprint for Neuroscience Research; and by the McDonnell Center for Systems Neuroscience at Washington University.

---

**Table S1:** Parameters used in backward reconstruction

| Variable | Annotation | HCP<br>(B=3T) | Mouse<br>(B=9.4T) |
| --- | --- | --- | --- |
| $\vartheta_0$ | Frequency offset at the outer surface<br>of the magnetized vessel for fully deoxygenated blood | 80.6 [2] | 252.5 [2] |
| $\rho$ | Oxygen extraction fraction | 0.8 | 0.8 |
| $\epsilon$ | Ratio of intra- and extravascular signal | 0.5 | 0.5 |
| $r_0$ | Slope of the relation between<br>intravascular relaxation rate and oxygen saturation | 167 [4] | 458 [1] |
| TE | Time of echo (ms) | 30 | 40 |
| $\tau$ | haemodynamic transit time | 0.98 | 0.98 |
| $\alpha$ | Grubbs exponent | 0.32 | 0.32 |
| $\kappa$ | Rate of signal decay | 0.65 | 0.65 |
| $\gamma$ | Rate of flow-dependent elimination | 0.41 | 0.41 |
| $\eta$ | Neuronal efficacy | 0.54 | 0.54 |
| $V_0$ | Resting blood volume fraction | 0.02 | 0.02 |

**Table S2:**
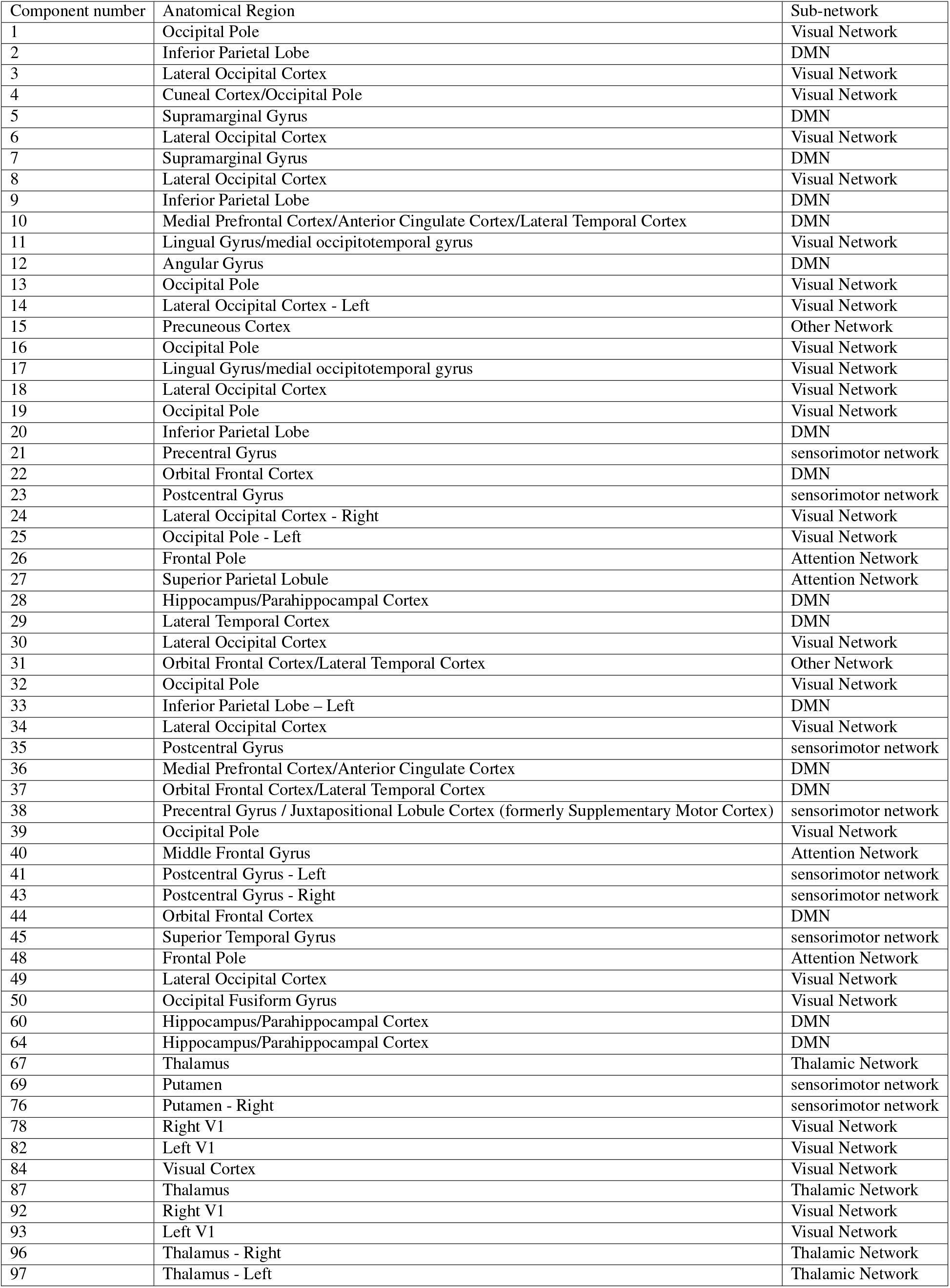
Annotated components for HCP dataset. First column is the index number of pre-selected IC components. Second column is the manually registered anatomical region. Third column is the assigned sub-network. DMN: default mode network

| Component number | Anatomical Region | Sub-network |
| --- | --- | --- |
| 1 | Occipital Pole | Visual Network |
| 2 | Inferior Parietal Lobe | DMN |
| 3 | Lateral Occipital Cortex | Visual Network |
| 4 | Cuneal Cortex/Occipital Pole | Visual Network |
| 5 | Supramarginal Gyrus | DMN |
| 6 | Lateral Occipital Cortex | Visual Network |
| 7 | Supramarginal Gyrus | DMN |
| 8 | Lateral Occipital Cortex | Visual Network |
| 9 | Inferior Parietal Lobe | DMN |
| 10 | Medial Prefrontal Cortex/Anterior Cingulate Cortex/Lateral Temporal Cortex | DMN |
| 11 | Lingual Gyrus/medial occipitotemporal gyrus | Visual Network |
| 12 | Angular Gyrus | DMN |
| 13 | Occipital Pole | Visual Network |
| 14 | Lateral Occipital Cortex - Left | Visual Network |
| 15 | Precuneous Cortex | Other Network |
| 16 | Occipital Pole | Visual Network |
| 17 | Lingual Gyrus/medial occipitotemporal gyrus | Visual Network |
| 18 | Lateral Occipital Cortex | Visual Network |
| 19 | Occipital Pole | Visual Network |
| 20 | Inferior Parietal Lobe | DMN |
| 21 | Precentral Gyrus | sensorimotor network |
| 22 | Orbital Frontal Cortex | DMN |
| 23 | Postcentral Gyrus | sensorimotor network |
| 24 | Lateral Occipital Cortex - Right | Visual Network |
| 25 | Occipital Pole - Left | Visual Network |
| 26 | Frontal Pole | Attention Network |
| 27 | Superior Parietal Lobule | Attention Network |
| 28 | Hippocampus/Parahippocampal Cortex | DMN |
| 29 | Lateral Temporal Cortex | DMN |
| 30 | Lateral Occipital Cortex | Visual Network |
| 31 | Orbital Frontal Cortex/Lateral Temporal Cortex | Other Network |
| 32 | Occipital Pole | Visual Network |
| 33 | Inferior Parietal Lobe – Left | DMN |
| 34 | Lateral Occipital Cortex | Visual Network |
| 35 | Postcentral Gyrus | sensorimotor network |
| 36 | Medial Prefrontal Cortex/Anterior Cingulate Cortex | DMN |
| 37 | Orbital Frontal Cortex/Lateral Temporal Cortex | DMN |
| 38 | Precentral Gyrus / Juxtapositional Lobule Cortex (formerly Supplementary Motor Cortex) | sensorimotor network |
| 39 | Occipital Pole | Visual Network |
| 40 | Middle Frontal Gyrus | Attention Network |
| 41 | Postcentral Gyrus - Left | sensorimotor network |
| 43 | Postcentral Gyrus - Right | sensorimotor network |
| 44 | Orbital Frontal Cortex | DMN |
| 45 | Superior Temporal Gyrus | sensorimotor network |
| 48 | Frontal Pole | Attention Network |
| 49 | Lateral Occipital Cortex | Visual Network |
| 50 | Occipital Fusiform Gyrus | Visual Network |
| 60 | Hippocampus/Parahippocampal Cortex | DMN |
| 64 | Hippocampus/Parahippocampal Cortex | DMN |
| 67 | Thalamus | Thalamic Network |
| 69 | Putamen | sensorimotor network |
| 76 | Putamen - Right | sensorimotor network |
| 78 | Right V1 | Visual Network |
| 82 | Left V1 | Visual Network |
| 84 | Visual Cortex | Visual Network |
| 87 | Thalamus | Thalamic Network |
| 92 | Right V1 | Visual Network |
| 93 | Left V1 | Visual Network |
| 96 | Thalamus - Right | Thalamic Network |
| 97 | Thalamus - Left | Thalamic Network |

**Table S3:** Annotated IC maps for the Bukhari mouse dataset. First column is the index of pre-selected IC components. Second column is the manually registered anatomical region. Third column is the corresponding anatomical locations in the Swanson database [3]. SSp = Primary somatosensory area, SSs = Supplemental somatosensory area, MO = Somatomotor areas, ACAd(v) = Anterior cingulate area dorsal (ventral), RSPd(v) = Retrosplenial region dorsal (ventral), CEA = Central amygdalar nucleus, MEA = Medial amygdalar nucleus, AAA = Anterior amygdalar area, BMA = Basomedial amygdalar nucleus, PAA = Piriform-amygdalar area, IA = Intercalated amygdalar nuclei, AAA = Anterior amygdalar area, BLA = Basolateral amygdalar nucleus, LA = Lateral amygdalar nucleus, PA = Posterior amygdalar nucleus, ECT = Ectorhinal area, PERI = Perirhinal area, CP = Caudoputamen, AI = Agranular insular, PIR = Piriform area, GP = Globus pallidus, NA: no corresponding anatomical locations in the database

| IC number | Full Name | Swanson region (Abbreviations) | Subnetwork |
| --- | --- | --- | --- |
| 1 | Primary Somatosensory Cortex (both side) | SSp | Lateral cortical network |
| 2 | Secondary Sematosensory Cortex (right) | SSs | Lateral cortical network |
| 3 | Secondary Sematosensory Cortex (left) | SSs | Lateral cortical network |
| 4 | Primary Somatosensory Cortex (right) | SSp | Lateral cortical network |
| 5 | Primary Somatosensory Cortex (left) | SSp | Lateral cortical network |
| 6 | Motor Cortex (right) | MO | Lateral cortical network |
| 7 | Motor Cortex (left) | MO | Lateral cortical network |
| 8 | Anterior Cingulate Cortex, dorsal | ACAd | Default mode network |
| 9 | Cingulate Cortex/ Retrosplenial Cortex | ACAv/ACAd/RSPd/RSPv | Default mode network |
| 10 | Amygdalar (right) | CEA/MEA/AAA/BMA/PAA/IA/AAA/BLA/LA/PA | Associative cortical network |
| 11 | Amygdalar (left) | CEA/MEA/AAA/BMA/PAA/IA/AAA/BLA/LA/PA | Associative cortical network |
| 12 | Ectorhinal Cortex (left) | ECT | Associative cortical network |
| 13 | Ectorhinal Cortex (right) | ECT | Associative cortical network |
| 14 | Perirhinal Area (right) | PERI | Associative cortical network |
| 15 | Caudoputamen (right) | CP | Subcortical network |
| 16 | Caudoputamen (left) | CP | Subcortical network |
| 17 | Insular (left) | AI | Associative cortical network |
| 18 | Insular (right) | AI | Associative cortical network |
| 19 | Insular (left) | AI | Associative cortical network |
| 20 | Insular (right) | AI | Associative cortical network |
| 21 | Perirhinal Area (right) | PERI | Associative cortical network |
| 22 | Perirhinal Area (left) | PERI | Associative cortical network |
| 23 | Piriform Cortex (left) | PIR | Associative cortical network |
| 24 | Piriform Cortex (right) | PIR | Associative cortical network |
| 25 | Globus Pallidus (right) | GP | Subcortical network |
| 26 | Globus Pallidus (left) | GP | Subcortical network |
| 27 | Thalamus, dorsal (left) | NA | Thalamic network |
| 28 | Thalamus, dorsal (right) | NA | Thalamic network |
| 29 | Thalamus, ventral (left) | NA | Thalamic network |
| 30 | Thalamus, ventral (right) | NA | Thalamic network |

**Figure S1:**
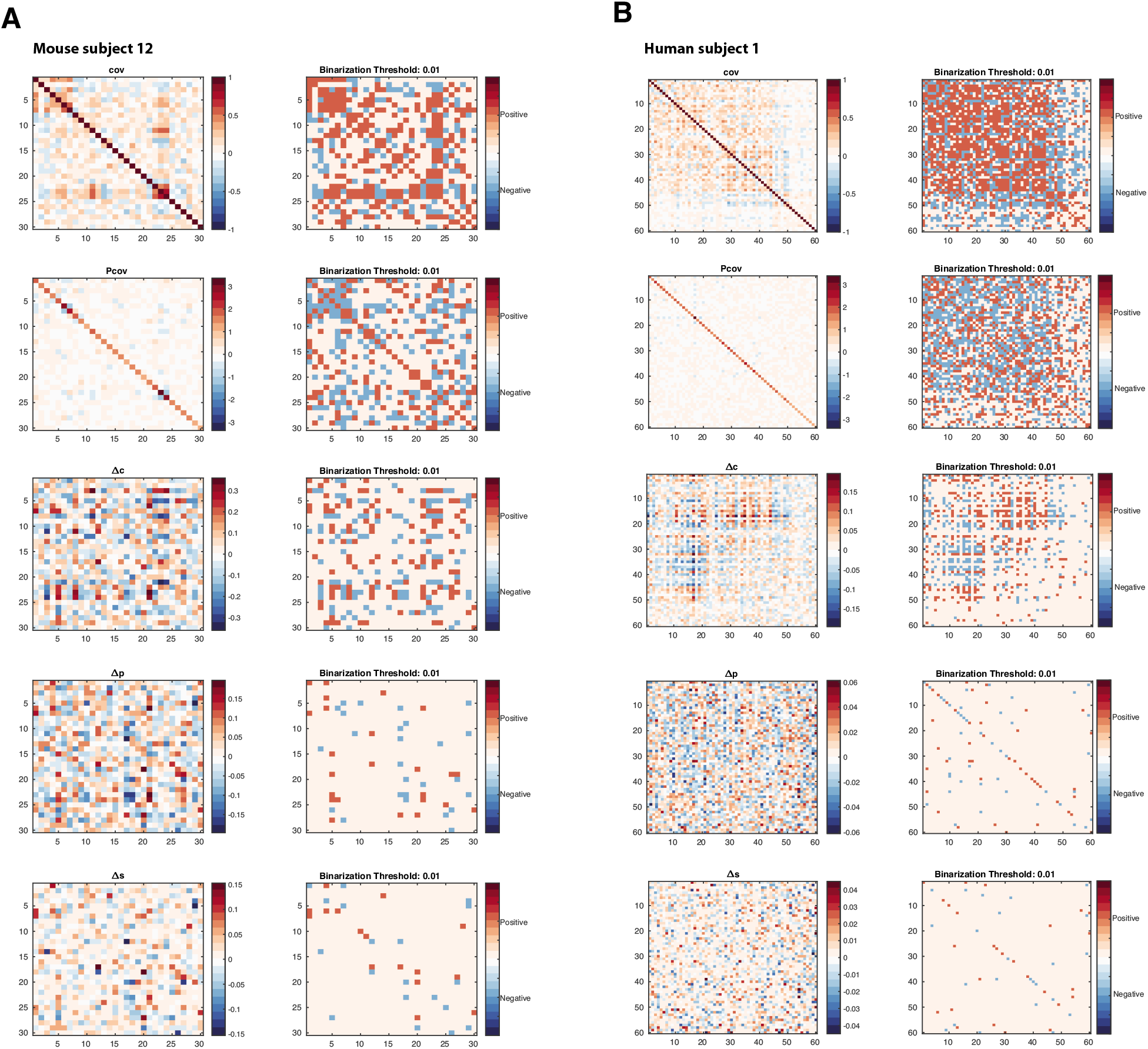
Differential covariance based methods produce sparse and directed FC. Example covariance matrix (cov), precision matrix (Pcov), Δ*c*, Δ*p* and Δ*s* and their binarized matrices (right) of one mouse subject (A) and one human subject (B).

**Figure S2:**
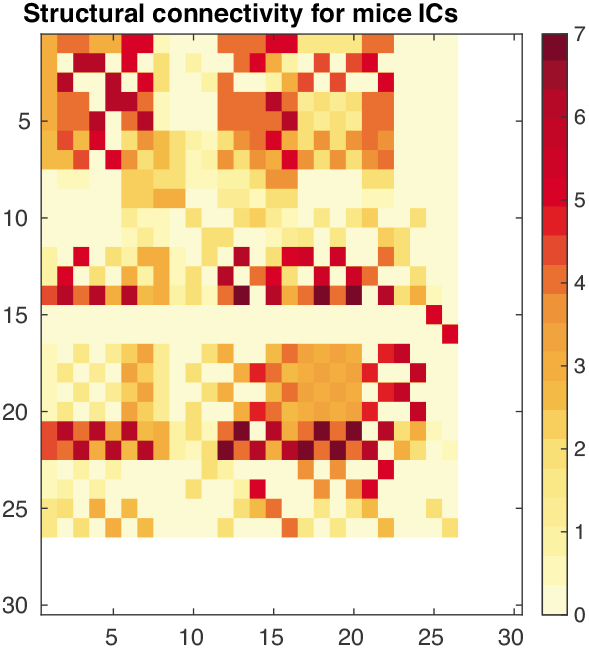
The structural connectivity matrix for mouse.

**Figure S3:**
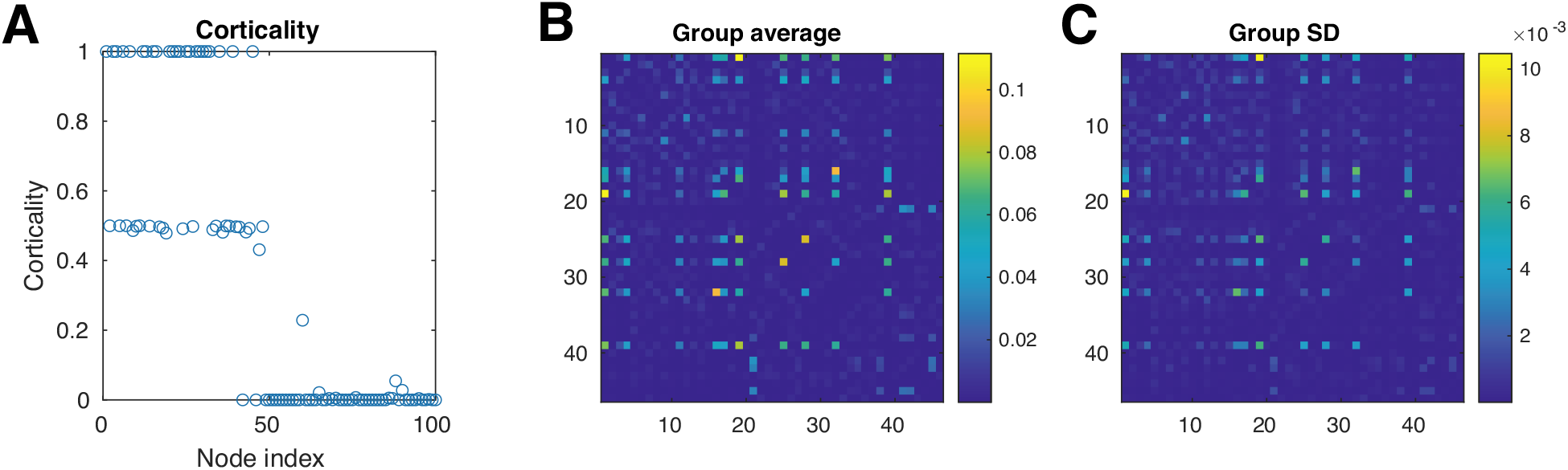
Individual level dMRI statistics. (A) Corticality was defined as the proportion of cortical voxels within each IC. Since dMRI measurements are only available for cortical surface voxels. Our analysis was restricted to the first 46 ICs with corticality greater than 40%. (B) Average of the dMRI matrices across the entire 998 subjects. (C) Standard deviation.

**Figure S4:**
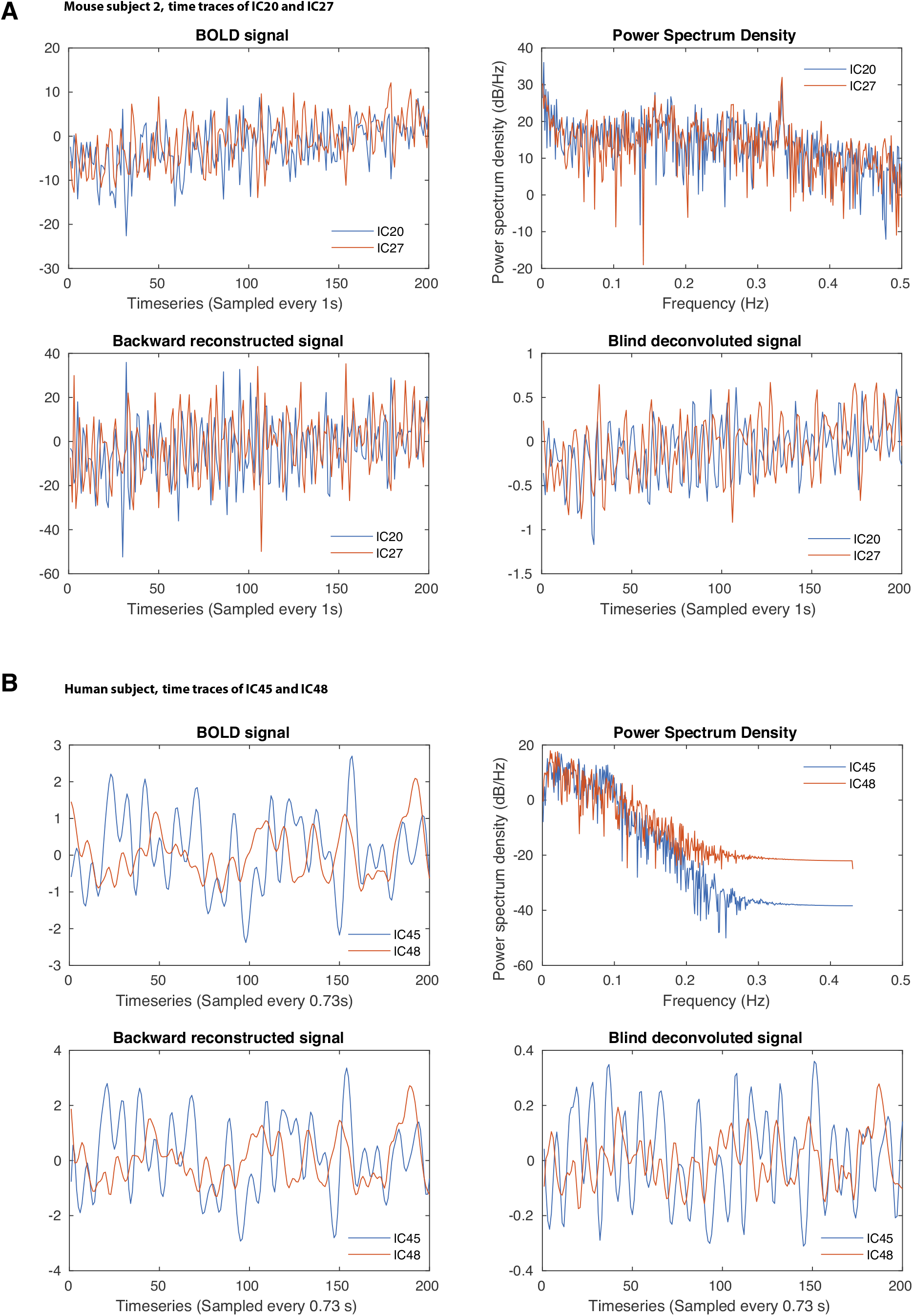
Two example time traces from one mouse subject (A) and one human subject (B) Upper left: haemodynamic signal after dual reg7ression; Upper right: power spectrum density of the haemodynamic signal; Lower left: backward reconstructed signal; Lower right: blind deconvoluted signal

**Figure S5:**
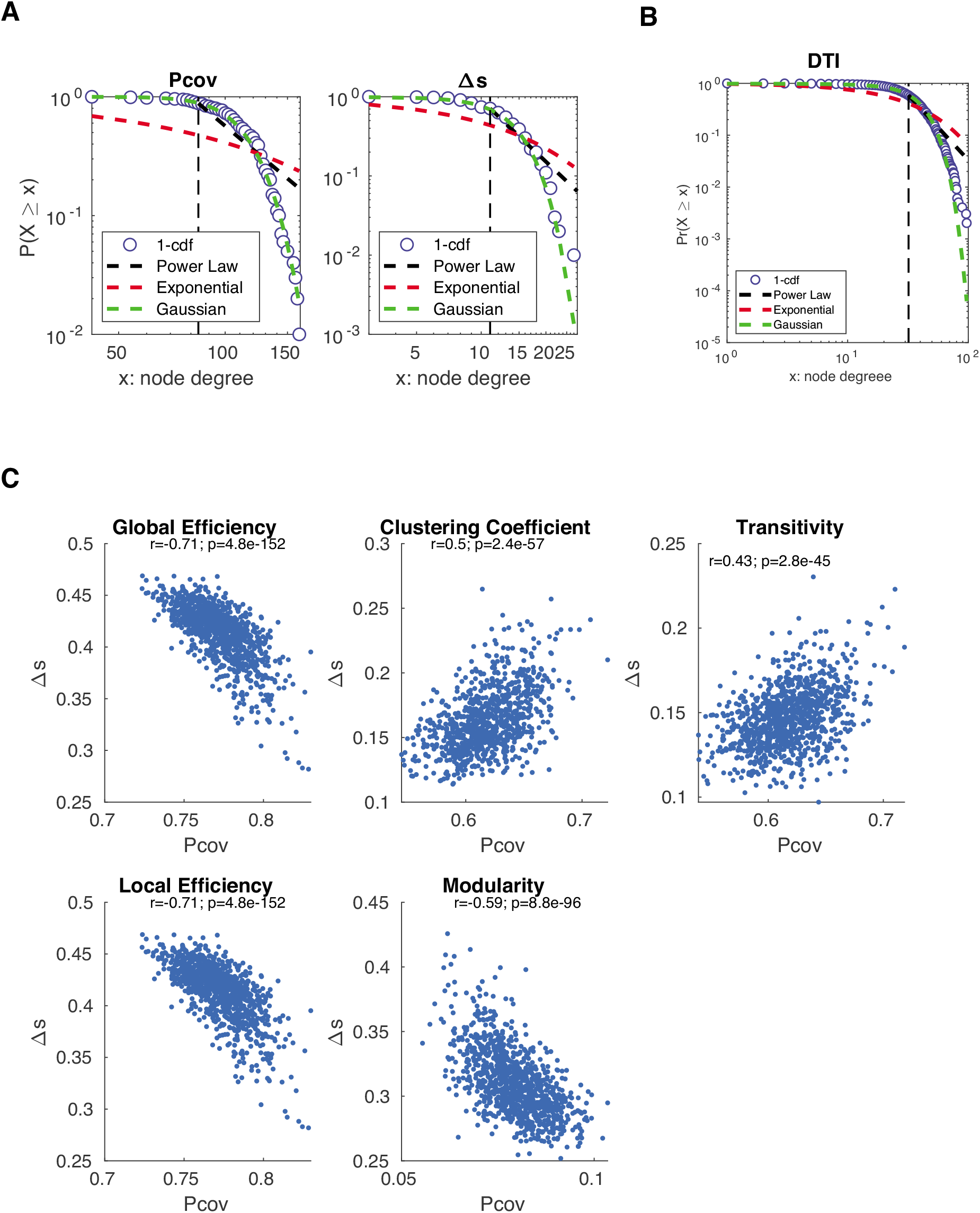
Network topology analysis of FC. (A) Cumulative density function of network degree distribution of Pcov-FC and Δ*s*-FC from one HCP subject. (B) Network degree distribution of structural connectivity matrix from diffusion tensor imaging (DTI). (C) Scatterplots of global efficiency, local efficiency, clustering coefficient, transitivity and modularity of Δ*s*-FC versus that of Pcov-FC.

1 Note that the acronym dCov-FC refers to functional connectivity revealed by differential covariance. This should not be confused with the distance covariance based upon pairwise distances between random variables https://en.wikipedia.org/wiki/Distance_correlation

## References

[1] Farras Abdelnour, Michael Dayan, Orrin Devinsky, Thomas Thesen, and Ashish Raj. Functional brain connectivity is predictable from anatomic network’s laplacian eigen-structure. NeuroImage, 172:728–739, 2018.

[2] Farras Abdelnour, Henning U Voss, and Ashish Raj. Network diffusion accurately models the relationship between structural and functional brain connectivity networks. Neuroimage, 90:335–347, 2014.

[3] Andrea Avena-Koenigsberger, Bratislav Misic, and Olaf Sporns. Communication dynamics in complex brain networks. Nature Reviews Neuroscience, 19(1):17, 2018.

[4] Qasim Bukhari, Aileen Schroeter, and Markus Rudin. Increasing isoflurane dose reduces homotopic correlation and functional segregation of brain networks in mice as revealed by resting-state fmri. Scientific reports, 8(1):10591, 2018.

[5] Emmanuel J Candès, Xiaodong Li, Yi Ma, and John Wright. Robust principal component analysis? Journal of the ACM (JACM), 58(3):1–37, 2011.

[6] Yusi Chen, Burke Q Rosen, and Terrence J Sejnowski. Dynamical differential covariance recovers directional network structure in multiscale neural systems. bioRxiv, 2021.

[7] Jessica R Cohen and Mark D’Esposito. The segregation and integration of distinct brain networks and their relationship to cognition. Journal of Neuroscience, 36(48):12083–12094, 2016.

[8] David Roxbee Cox and Nanny Wermuth. Multivariate dependencies: Models, analysis and interpretation. Chapman and Hall/CRC, 2014.

[9] Bradley Efron and Robert Tibshirani. Bootstrap methods for standard errors, confidence intervals, and other measures of statistical accuracy. Statistical science, pages 54–75, 1986.

[10] Nicola Filippini, Bradley J MacIntosh, Morgan G Hough, Guy M Goodwin, Giovanni B Frisoni, Stephen M Smith, Paul M Matthews, Christian F Beckmann, and Clare E Mackay. Distinct patterns of brain activity in young carriers of the apoe-*ε*4 allele. Proceedings of the National Academy of Sciences, 106(17):7209–7214, 2009.

[11] Stefan Frässle, Ekaterina I Lomakina, Adeel Razi, Karl J Friston, Joachim M Buhmann, and Klaas E Stephan. Regression dcm for fmri. Neuroimage, 155:406–421, 2017.

[12] Karl J Friston, Lee Harrison, and Will Penny. Dynamic causal modelling. Neuroimage, 19(4):1273–1302, 2003.

[13] Karl J Friston, Andrea Mechelli, Robert Turner, and Cathy J Price. Nonlinear responses in fmri: the balloon model, volterra kernels, and other hemodynamics. NeuroImage, 12(4):466–477, 2000.

[14] Matthew F Glasser, Timothy S Coalson, Emma C Robinson, Carl D Hacker, John Harwell, Essa Yacoub, Kamil Ugurbil, Jesper Andersson, Christian F Beckmann, Mark Jenkinson, et al. A multi-modal parcellation of human cerebral cortex. Nature, 536(7615):171, 2016.

[15] Matthew F Glasser, Stamatios N Sotiropoulos, J Anthony Wilson, Timothy S Coalson, Bruce Fischl, Jesper L Andersson, Junqian Xu, Saad Jbabdi, Matthew Webster, Jonathan R Polimeni, et al. The minimal preprocessing pipelines for the human connectome project. Neuroimage, 80:105–124, 2013.

[16] Joanes Grandjean, Valerio Zerbi, Joshua Henk Balsters, Nicole Wenderoth, and Markus Rudin. Structural basis of large-scale functional connectivity in the mouse. Journal of Neuroscience, 37(34):8092–8101, 2017.

[17] Ludovica Griffanti, Gholamreza Salimi-Khorshidi, Christian F Beckmann, Edward J Auerbach, Gwenaëlle Douaud, Claire E Sexton, Enikő Zsoldos, Klaus P Ebmeier, Nicola Filippini, Clare E Mackay, et al. Ica-based artefact removal and accelerated fmri acquisition for improved resting state network imaging. Neuroimage, 95:232–247, 2014.

[18] Patric Hagmann, Leila Cammoun, Xavier Gigandet, Reto Meuli, Christopher J Honey, Van J Wedeen, and Olaf Sporns. Mapping the structural core of human cerebral cortex. PLoS biology, 6(7):e159, 2008.

[19] CJ Honey, O Sporns, Leila Cammoun, Xavier Gigandet, Jean-Philippe Thiran, Reto Meuli, and Patric Hagmann. Predicting human resting-state functional connectivity from structural connectivity. Proceedings of the National Academy of Sciences, 106(6):2035–2040, 2009.

[20] Li-Chuan Huang, Ping-An Wu, Shinn-Zong Lin, Cheng-Yoong Pang, and Shin-Yuan Chen. Graph theory and network topological metrics may be the potential biomarker in parkinson’s disease. Journal of Clinical Neuroscience, 68:235–242, 2019.

[21] Derek K Jones, Thomas R Knösche, and Robert Turner. White matter integrity, fiber count, and other fallacies: the do’s and don’ts of diffusion mri. Neuroimage, 73:239–254, 2013.

[22] Ildar Khalidov, Jalal Fadili, François Lazeyras, Dimitri Van De Ville, and Michael Unser. Activelets: Wavelets for sparse representation of hemodynamic responses. Signal processing, 91(12):2810–2821, 2011.

[23] Tiger W. Lin, Yusi Chen, Qasim Bukhari, Giri P. Krishnan, Maxim Bazhenov, and Terrence J. Sejnowski. Differential covariance: A new method to estimate functional connectivity in fmri. Neural Computation, 0(0):1–33, 0. PMID: 32946714.

[24] Tiger W Lin, Anup Das, Giri P Krishnan, Maxim Bazhenov, and Terrence J Sejnowski. Differential covariance: A new class of methods to estimate sparse connectivity from neural recordings. Neural computation, 29(10):2581–2632, 2017.

[25] Thomas T Liu. Noise contributions to the fmri signal: An overview. NeuroImage, 143:141–151, 2016.

[26] Scott Marek, Kai Hwang, William Foran, Michael N Hallquist, and Beatriz Luna. The contribution of network organization and integration to the development of cognitive control. PLoS biology, 13(12):e1002328, 2015.

[27] Arnaud Messé, David Rudrauf, Alain Giron, and Guillaume Marrelec. Predicting functional connectivity from structural connectivity via computational models using mri: an extensive comparison study. NeuroImage, 111:65–75, 2015.

[28] Alican Nalci, Bhaskar D Rao, and Thomas T Liu. Nuisance effects and the limitations of nuisance regression in dynamic functional connectivity fmri. NeuroImage, 184:1005–1031, 2019.

[29] Hae-Jeong Park, Karl J Friston, Chongwon Pae, Bumhee Park, and Adeel Razi. Dynamic effective connectivity in resting state fmri. Neuroimage, 180:594–608, 2018.

[30] Andrew T Reid, Drew B Headley, Ravi D Mill, Ruben Sanchez-Romero, Lucina Q Uddin, Daniele Marinazzo, Daniel J Lurie, Pedro A Valdés-Sosa, Stephen José Hanson, Bharat B Biswal, et al. Advancing functional connectivity research from association to causation. Nature neuroscience, 22(11):1751–1760, 2019.

[31] Emma C Robinson, Saad Jbabdi, Matthew F Glasser, Jesper Andersson, Gregory C Burgess, Michael P Harms, Stephen M Smith, David C Van Essen, and Mark Jenkinson. Msm: a new flexible framework for multimodal surface matching. Neuroimage, 100:414–426, 2014.

[32] Burke Q Rosen and Eric Halgren. A whole-cortex probabilistic diffusion tractography connectome. Eneuro, 8(1), 2021.

[33] Mikail Rubinov and Olaf Sporns. Complex network measures of brain connectivity: uses and interpretations. Neuroimage, 52(3):1059–1069, 2010.

[34] Gholamreza Salimi-Khorshidi, Gwenaëlle Douaud, Christian F Beckmann, Matthew F Glasser, Ludovica Griffanti, and Stephen M Smith. Automatic denoising of functional mri data: combining independent component analysis and hierarchical fusion of classifiers. Neuroimage, 90:449–468, 2014.

[35] Emiliano Santarnecchi, Giulia Galli, Nicola Riccardo Polizzotto, Alessandro Rossi, and Simone Rossi. Efficiency of weak brain connections support general cognitive functioning. Human brain mapping, 35(9):4566–4582, 2014.

[36] Stephen M Smith, Christian F Beckmann, Jesper Andersson, Edward J Auerbach, Janine Bijsterbosch, Gwenaëlle Douaud, Eugene Duff, David A Feinberg, Ludovica Griffanti, Michael P Harms, et al. Resting-state fmri in the human connectome project. Neuroimage, 80:144–168, 2013.

[37] Stephen M Smith, Aapo Hyvärinen, Gaél Varoquaux, Karla L Miller, and Christian F Beckmann. Group-pca for very large fmri datasets. Neuroimage, 101:738–749, 2014.

[38] Stephen M Smith, Karla L Miller, Gholamreza Salimi-Khorshidi, Matthew Webster, Christian F Beckmann, Thomas E Nichols, Joseph D Ramsey, and Mark W Woolrich. Network modelling methods for fmri. Neuroimage, 54(2):875–891, 2011.

[39] Stephen M Smith, Thomas E Nichols, Diego Vidaurre, Anderson M Winkler, Timothy EJ Behrens, Matthew F Glasser, Kamil Ugurbil, Deanna M Barch, David C Van Essen, and Karla L Miller. A positive-negative mode of population covariation links brain connectivity, demographics and behavior. Nature neuroscience, 18(11):1565, 2015.

[40] Olaf Sporns. Network attributes for segregation and integration in the human brain. Current opinion in neurobiology, 23(2):162–171, 2013.

[41] Klaas Enno Stephan, Lars Kasper, Lee M Harrison, Jean Daunizeau, Hanneke EM den Ouden, Michael Breakspear, and Karl J Friston. Nonlinear dynamic causal models for fmri. Neuroimage, 42(2):649–662, 2008.

[42] Klaas Enno Stephan, Nikolaus Weiskopf, Peter M Drysdale, Peter A Robinson, and Karl J Friston. Comparing hemodynamic models with dcm. Neuroimage, 38(3):387–401, 2007.

[43] Ian H Stevenson, James M Rebesco, Lee E Miller, and Konrad P Körding. Inferring functional connections between neurons. Current opinion in neurobiology, 18(6):582–588, 2008.

[44] Larry W Swanson, Joel D Hahn, Lucas GS Jeub, Santo Fortunato, and Olaf Sporns. Subsystem organization of axonal connections within and between the right and left cerebral cortex and cerebral nuclei (endbrain). Proceedings of the National Academy of Sciences, 115(29):E6910–E6919, 2018.

[45] David C Van Essen, Stephen M Smith, Deanna M Barch, Timothy EJ Behrens, Essa Yacoub, Kamil Ugurbil, Wu-Minn HCP Consortium, et al. The wu-minn human connectome project: an overview. Neuroimage, 80:62–79, 2013.

[46] Justin L Vincent, Gaurav H Patel, Michael D Fox, Abraham Z Snyder, Justin T Baker, David C Van Essen, John M Zempel, Lawrence H Snyder, Maurizio Corbetta, and Marcus E Raichle. Intrinsic functional architecture in the anaesthetized monkey brain. Nature, 447(7140):83, 2007.

[47] Guo-Rong Wu, Wei Liao, Sebastiano Stramaglia, Ju-Rong Ding, Huafu Chen, and Daniele Marinazzo. A blind deconvolution approach to recover effective connectivity brain networks from resting state fmri data. Medical image analysis, 17(3):365–374, 2013.

[48] Dimitri Yatsenko, Krešimir Josić, Alexander S Ecker, Emmanouil Froudarakis, R James Cotton, and Andreas S Tolias. Improved estimation and interpretation of correlations in neural circuits. PLoS computational biology, 11(3):e1004083, 2015.

## References

[1] Sang-Pil Lee, Afonso C Silva, Kamil Ugurbil, and Seong-Gi Kim. Diffusion-weighted spinecho fmri at 9.4 t: microvascular/tissue contribution to bold signal changes. Magnetic Resonance in Medicine: An Official Journal of the International Society for Magnetic Resonance in Medicine, 42(5):919–928, 1999.

[2] S Ogawa, RS Menon, David W Tank, SG Kim, H Merkle, JM Ellermann, and K Ugurbil. Functional brain mapping by blood oxygenation level-dependent contrast magnetic resonance imaging. a comparison of signal characteristics with a biophysical model. Biophysical journal, 64(3):803–812, 1993.

[3] Larry W Swanson, Joel D Hahn, Lucas GS Jeub, Santo Fortunato, and Olaf Sporns. Subsystem organization of axonal connections within and between the right and left cerebral cortex and cerebral nuclei (endbrain). Proceedings of the National Academy of Sciences, 115(29):E6910–E6919, 2018.

[4] Jason M Zhao, Chekesha S Clingman, M Johanna Närväinen, Risto A Kauppinen, and Peter CM van Zijl. Oxygenation and hematocrit dependence of transverse relaxation rates of blood at 3t. Magnetic Resonance in Medicine: An Official Journal of the International Society for Magnetic Resonance in Medicine, 58(3):592–597, 2007.

